# Fluoxetine Hydrochloride Treatment Influences Site-Specific ADAR Editing and Transcriptome Regulation in Arid1b +/- Mice

**DOI:** 10.64898/2026.09.24.753309

**Authors:** Ayesha Tariq, Helen Piontkivska

## Abstract

Autism Spectrum Disorder (ASD) is a neurodevelopmental disorder characterized by repetitive behavior and impaired social interactions. Recent reports from human postmortem brain samples of ASD show transcriptome-wide dysregulations, including changes in differential gene expression and alternative splicing. An important mechanism regulating transcriptome function is the editing of double-stranded RNA by adenosine deaminases acting on RNA (ADARs), which play regulatory roles in neurodevelopment and innate immunity. We explore the ADAR editing changes associated with the use of Selective Serotonin Reuptake Inhibitors (SSRIs) in Arid1b haploinsufficient mouse models. Our results show dynamic changes in site-specific editing rates in autistic mice. We further demonstrate the transcriptome-wide effects of differential ADAR editing in terms of changes in microRNA-mRNA interactions. Moreover, our findings highlight a site-specific change in editing rates for evolutionarily conserved sites in mammals and ASD associated risk genes, such as Gria2. Our analysis of significant biological processes shows the association of differentially edited genes with processes vital for neurotransmission. Overall, our findings imply that the use of SSRIs can induce distinct ADAR editing signatures, potentially contributing to ASD pathology.

## Introduction

Autism spectrum disorder (ASD) is a polygenic trait and one of the most prevalent neurodevelopmental disorders, affecting approximately 1 in every 44 children in the US alone (Maenner, 2023; Rylaarsdam and Guemez-Gamboa, 2019). ASD is commonly characterized by cognitive dysfunction, impaired social interaction, and repetitive behavior (Lord et al., 2020; Wang et al., 2023). Genetic risk factors associated with ASD include small genetic variants, rare copy number variants, ASD-related genetic syndromes, and chromosomal abnormalities (Choi and An, 2021; Genovese and Butler, 2023; Rylaarsdam and Guemez-Gamboa, 2019). Additionally, abnormal expression and nucleotide variations in microRNAs (miRNAs) may contribute by affecting the expression of ASD-associated risk genes (Garrido-Torres et al., 2024; Li et al., 2022; Wong et al., 2022). Among environmental factors, childhood infections and maternal immune activation (MIA) due to in utero infections lead to inflammatory perturbations and can potentially contribute to increased risk of neurodevelopmental disorders, including ASD (Atladóttir et al., 2010; Han et al., 2021; Nudel et al., 2022; Sabourin et al., 2019). Innate immune activation by polyinosinic polycytidylic acid (poly I:C) induction in rodents also supports the association between increased gestational inflammation and dysregulation of neurodevelopment (Gzieło et al., 2023; Haddad et al., 2020; Lan et al., 2023; Tartaglione et al., 2022). Increased inflammation due to MIA, in turn leading to dysregulation of RNA editing by ADARs (<u>A</u>denosine <u>D</u>eaminases <u>A</u>cting on <u>R</u>NA), may serve as one causative factor in dysregulation of neurodevelopment (Tsivion-Visbord et al., 2020; Wales-McGrath et al., 2023), although further validation work is needed.

Editing of double-stranded (ds) RNA by ADARs, also known as A-to-I editing or ADAR editing, is one of the most prevalent types of RNA editing in metazoans, and is a key regulator of neural development, plasticity, and aging (Gallo et al., 2017; Hwang et al., 2016). RNA editing by ADARs, through deamination of adenosines (A) to inosines (I), which are read as guanosines (G) by the translational machinery, contributes to the diversity of key neuronal receptor proteins (Behm and Öhman, 2016; Huntley et al., 2016; Walkley and Li, 2017). Three members of the ADAR gene family are Adar1 (Adar), Adar2 (Adarb1), and Adar3 (Adarb2); however, to date, catalytic activity has been documented only for Adar1 and Adar2 (Bass, 2002; Yang et al., 2021). While Adar2 has critical targets in the brain (Cha et al., 1994; Tanaka et al., 2000), Adar1p150, a long isoform of Adar1, is an interferon-stimulated gene (ISG), mediating the role of ADAR editing in both innate immunity and inflammatory processes, including during infections, autoimmune diseases, and neurological disorders (Piontkivska et al., 2019; Slotkin and Nishikura, 2013; Tariq and Piontkivska, 2024; Wales-McGrath et al., 2023; Yang et al., 2021). Moreover, reports from human postmortem brain tissues have identified differential transcriptome-wide ADAR editing in ASD, where tissue-dependent increase or decrease in site-specific editing was reported (Eran et al., 2013; Tran et al., 2019). Neuroinflammation, synaptic dysfunction, and abnormalities in neurotransmission in autism and related phenotypes (Eissa et al., 2020; Rubenstein and Merzenich, 2003; Sztainberg and Zoghbi, 2016; Than et al., 2023), and the role of ADAR editing in neurodevelopment, synaptic plasticity, and inflammation (Gallo et al., 2017; Hwang et al., 2016; Yang et al., 2021) make its RNA editing function a critical pathway in ASD.

The 5-hydroxytryptamine subtype 2C receptor (Htr2c) is the only member of the serotonin receptor family subjected to ADAR editing (Slotkin and Nishikura, 2013). In Htr2c mRNA, five conserved positions (A, B, C, D, and E) are edited, where both Adar1 and Adar2 have selective and overlapping targets, leading to over 24 mRNA isoforms (Burns et al., 1997). Dysregulation of Htr2c editing, which leads to variations in mRNA isoforms, has been associated with anxiety, bipolar disorder, suicide, and depression (Dracheva et al., 2008; Gurevich et al., 2002; Iwamoto and Kato, 2003; Weissmann et al., 2016). While no approved therapies for ASD core symptoms are available, selective serotonin reuptake inhibitors (SSRIs) and treatments that target serotonergic activity are some of the therapeutic options to treat ASD-related anxiety and depression (Lee et al., 2022; Sclabassi et al., 2025). Fluoxetine hydrochloride, or fluoxetine, is a highly selective SSRI and a widely prescribed antidepressant that has been shown to alleviate some autism-related phenotypes (Kim et al., 2022; Wong et al., 2005, 1974). These drugs inhibit serotonin (5-hydroxytryptamine, 5-HT) reuptake by blocking the presynaptic serotonin transporter (SERT), thereby increasing extracellular serotonin availability (Fuller et al., 1991; Stahl, 1998). Although prescribed as therapeutics, SSRIs can also have adverse effects. Prenatal exposure to SSRIs, including fluoxetine, can block placental serotonin receptors, causing an increase in serotonin levels, leading to downstream consequences for fetal brain development, including an increased risk of ASD (Horackova et al., 2021; Man et al., 2015). Moreover, chronic fluoxetine treatment in adult mice leads to changes in A-to-I editing of Htr2c mRNA (Englander et al., 2005), while pregestational exposure in rats leads to changes in A-to-I editing of glutamate and serotonin receptors (Zaidan et al., 2018), suggesting off-target effects of SSRIs.

Given the above-described role of ADARs in neurodevelopment and transcriptome regulation, a comprehensive landscape of ADAR editing after SSRI exposure may provide additional insight into SSRIs’ mechanism of action and potential consequences as a therapeutic option for ASD. Thus, to explore the aforementioned dynamics of SSRI use and ADAR editing, we used publicly available deep-sequenced RNA-seq data from Arid1b haploinsufficient mouse models (Kim et al., 2022). Our results show distinct ADAR editing changes associated with autistic phenotype and SSRI use. We observed site-specific changes in ASD associated risk genes and changes in editing rates at evolutionarily conserved ADAR targets. Furthermore, we demonstrate the effect of editing on transcriptome regulation in terms of miRNA-mRNA interactions. Overall, our findings highlight differential ADAR editing associated with the use of SSRIs in ASD and provide a subset of editing signatures for future studies.

## Methods

### RNA-Sequencing Data

We used publicly available BioProject PRJNA839604 transcriptomics dataset from Kim et al. (2022) (GEO accession number GSE203343), where total RNA was extracted from the prefrontal cortex (PFC) of Arid1b haploinsufficient (Arid1b+/-) mice. Briefly, the experimental group consisted of 16 C57BL/6N male mice, including 8 wild-type (WT), and 8 Arid1b heterozygous (HT) mice. ARID1B is a component of the BAF chromatin remodeling complex, and its dysfunction causes autistic phenotypes such as decreased social communication and repetitive behavior, both in humans and mouse models (Jung et al., 2017). In each group, 4 mice were treated with fluoxetine hydrochloride (0.1 mg/ml in 0.2% saccharin) through mother’s milk, while the other 4 in each group were treated with 0.2% saccharin water, called the vehicle, from postnatal days 3-21 (P3-P21). After treatment with the drug and vehicle, total RNA was extracted from the PFC at P120, and deep-sequenced. HT and WT mice treated with fluoxetine or vehicle will be described as HT-drug, WT-drug, HT-veh, and WT-veh, respectively.

### Gene Expression Analysis

Analysis for differentially expressed genes (DEGs) was done using the DESeq2 R package (Love et al., 2014) with default parameters. For a direct comparison, normalized count values for all three ADAR genes (Adar1, Adar2, Adar3) were extracted using the function “normalized” in DESeq2. Normalized counts for ADAR genes were compared between HT and WT mice, using the R package ggstatplot’s (Patil, 2021) non-parametric approach for comparison between multiple groups. DEGs were filtered to include log2fold change > 0.58 or < -0.58, and BH-adjusted p-value ≤ 0.05.

### Quantification of ADAR Editing

The automated isoform diversity detector (AIDD) pipeline (Plonski et al., 2020) was used to map the reads to the reference genome GRCm38/mm10, perform variant calling, and predict ADAR editing sites. The editing rates were defined as the number of G reads for an A reference site or C reads for a T(U) reference site on the opposite strand, divided by the total number of reads mapped to that site. Mean editing per site was calculated as the average of editing rates for a single site across all samples for a specific condition. To ensure the inclusion of high confidence sites, edited sites with reference in the REDIportal, a comprehensive nonredundant database of A-to-I editing events (Picardi et al., 2017), were included. Putative ADAR edits were also filtered to remove known SNPs from dbSNP (Sherry et al., 2001) provided in REDIportal v2.0 (Mansi et al., 2021). Other filtering options required that the edited site had greater than or equal to 20 total reads aligned, the edited site is present in at least 50% of the samples, and that editing rates were greater than 0.01, or less than 0.99, and not between 0.49 and 0.51, to remove potential noise, and homozygous and heterozygous genomic variants, respectively. Differential editing inferences were drawn as: genomic sites edited in both HT and WT groups (shared sites), and sites only uniquely edited in either group (unique sites). To detect condition-specific changes in editing at shared sites, the difference in mean editing rate for shared sites was calculated with a 95% confidence interval (CI), based on the t-score for the difference in means. Use of CI limits over p-values allows filtering for not only significant hits but also provides an estimate of effect size for the observed differences (Hazra, 2017). Filtering criteria for unique sites included whether an edited site is edited in at least 50% of the samples, and that reads are not missing in any of the samples.

### Identification of miRNA Binding Targets

To compute miRNA targets for differentially edited sites, an approach similar to Roberts et al., (2018) was used. The list of differentially edited sites (shared and unique) was used to generate two data files for FASTA sequences of edited and unedited transcript versions. Each file contained a 201 base pairs (bp) long sequence for each edited site, comprising 100 bp upstream and downstream of the edited sites. Edited and unedited versions of FASTA sequence files were compared with the list of miRNAs for *Mus musculus*, from miRbase (Griffiths-Jones et al., 2006), using the RNA22 (v2) program (Miranda et al., 2006) with default parameters. The resulting miRNA-mRNA duplexes were filtered to include edited coordinates and p-value ≤ 0.05. The strength estimation of miRNA-mRNA heteroduplex was based on folding energy values. Folding energy or Gibbs free energy is the amount of energy gained when miRNA binds to its target (Ghoshal et al., 2015; Miranda et al., 2006). These values indicate the thermodynamic stability of the heteroduplex, where lower values indicate that more energy is required to break the heteroduplex, hence stronger binding (Ghoshal et al., 2015; Miranda et al., 2006). All the analyses were done using the GRCm38/mm10 mouse assembly (“GENCODE - Mouse Release M10,” 2024).

### Identification of Editing Changes in ASD Associated Risk Genes

Finally, significantly differentially edited gene targets were compared with the simons foundation autism research initiative (SFARI) Gene database (Abrahams et al., 2013) to filter for ASD-associated risk genes. SFARI Gene is a web-based database curated from peer-reviewed clinical and scientific knowledge about the molecular genetics and etiology of ASD. We cross-compared our data with the ASD associated risk gene list from mouse and human studies.

### Gene Ontology and Statistical Analysis

Gene ontology (GO) analysis for significant biological processes (BP) was performed using the R package clusterProfiler (Wu et al., 2021). Statistical analysis and visualization were performed in R version 4.3.1 (R Core Team, 2023). The citation list for additional R libraries used is provided in the supplemental files. Detailed results of GO analyses can be accessed by using gene lists provided in Supplemental Table 11 and R code files (See supplemental files).

## Results

### Fluoxetine Hydrochloride Treatment Impacts Global and Site-Specific ADAR Editing

All four experimental groups (HT-drug, HT-veh, WT-drug, WT-veh) were compared for the site-specific editing at unique and overlapping edited targets. While the majority of the edited sites are shared across groups, unique targets in each group, with the highest number in HT-veh, were observed (Figure 1-A, Supplemental Tables 2-5). The majority of the edited targets (overlapping and unique) are found in non-coding regulatory regions such as 3’UTRs, 5’UTRs, and introns (Figure 1-B), and the least number is present in non-coding RNAs (ncRNA) (Figure 1-D), while all the ADAR edits in exonic regions (Figure 1-C) are shared between all groups. The presence of overlapping and unique ADAR editing targets highlights the dynamic nature of ADAR editing, where editing events, depending upon the specific genomic coordinate, are dependent (unique) or independent (overlapping) of the treatment condition and genotype. Global editing between groups was compared as the total number of edited sites and mean editing rate per group. Although no statistically significant difference was found (Supplemental Figure 1-A), a comparison of site-specific editing rates for shared and unique targets shows that, compared to other treatment groups, fluoxetine treatment causes more distinct changes in the HT-drug group (Figure 1-E), indicating the incidence of ADAR editing on targets potentially associated with SRRIs’ mechanism of action.

**Figure 1.**
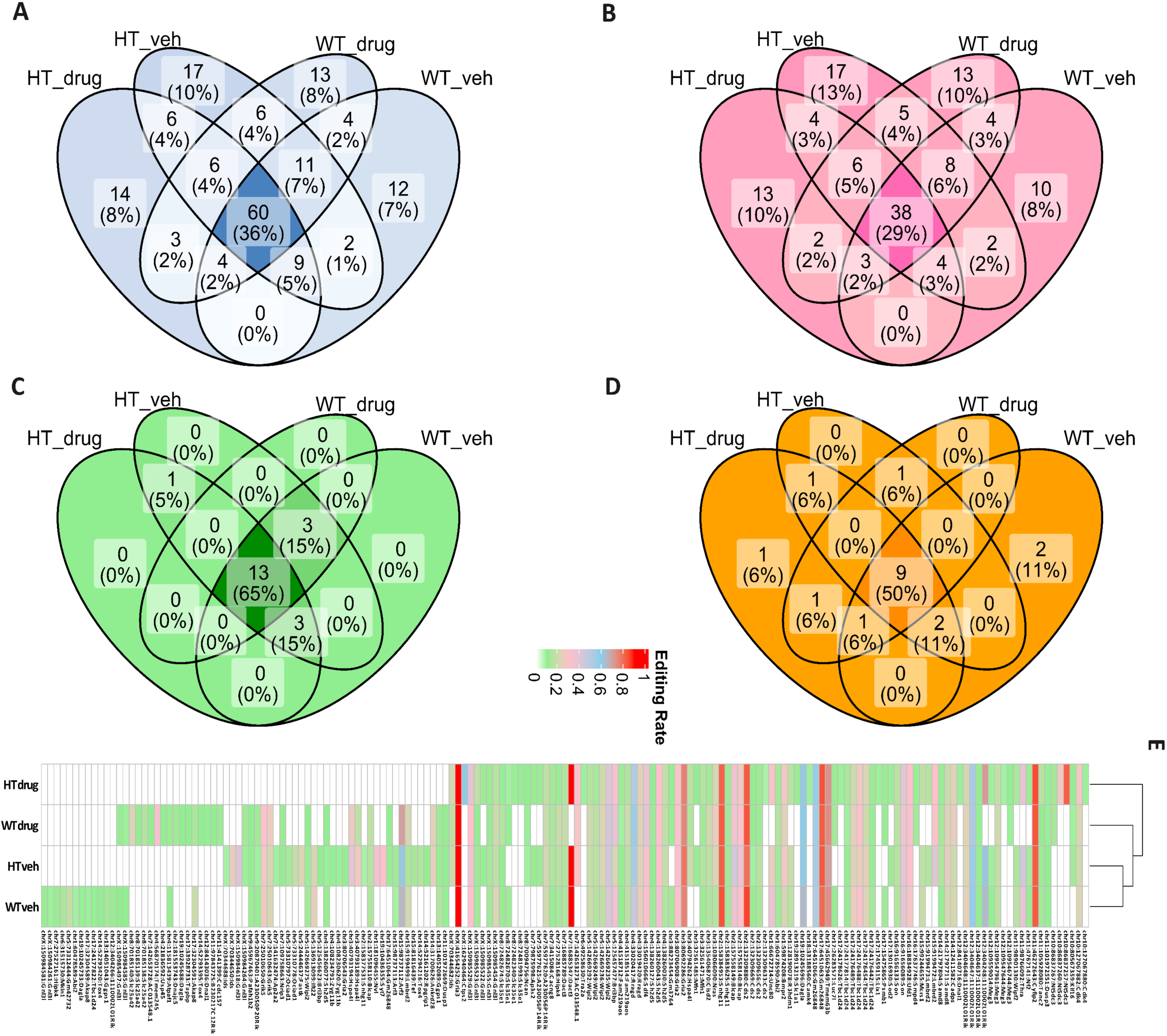
Transcriptome Wide ADAR Editing Landscape for Coding and Non-Coding Regions. Venn diagrams of region-specific ADAR-edited sites shown as **(A)** total ADAR-edited sites, edited sites in **(B)** non-coding regulatory regions such as 3’UTRs, 5’UTRs, and introns, **(C)** coding (exonic) regions, and **(D)** non-coding RNAs (ncRNAs). Set labels represent the treatment group and the number of edited sites in each category for each group. The number of overlapping and unique targets is represented by relative count and percentage of the edited sites in each category (i.e., region) for each group (i.e., treatment group). **(E)** A heatmap of site-specific editing rate for all treatment groups after exclusion of variants found in dbSNP and potential homozygous and heterozygous variants. Color key represents the magnitude of editing, where 0 indicates the absence of editing despite the presence of sequence reads for the respective genomic coordinate (Supplemental Table 12).

### ASD Associated Risk Genes and Evolutionarily Conserved ADAR Targets Experience Differential Editing in ASD Phenotype and Fluoxetine Treatment

Whether some editing changes could be exclusively attributed to fluoxetine treatment or the autistic phenotype, we compared site-specific editing rates between overlapping and unique targets. Experimental groups with fluoxetine or vehicle treatment were compared with each other. After the application of significance threshold cut-offs, we found 63 edited sites mapped to 45 genes shared between HT-drug and WT-drug samples, while there were 76 shared sites mapped to 46 genes between the HT-veh and WT-veh (Figure 2, Supplemental Tables 2 & 3). As a measure of effect size and significance, a 95% confidence interval (CI) around the difference in mean editing rate using t-critical score was calculated, and only intervals with non-zero upper and lower limits were included. As shown in Figure 2 (A), shared edited targets between vehicle- and drug-treated groups have significantly different editing rates, while differences in shared edited targets, e.g., Klf16, Nt5dc3 and Sdf4, which are only edited in fluoxetine treated groups, indicate the potential effect of fluoxetine treatment on ADAR editing.

**Figure 2.**
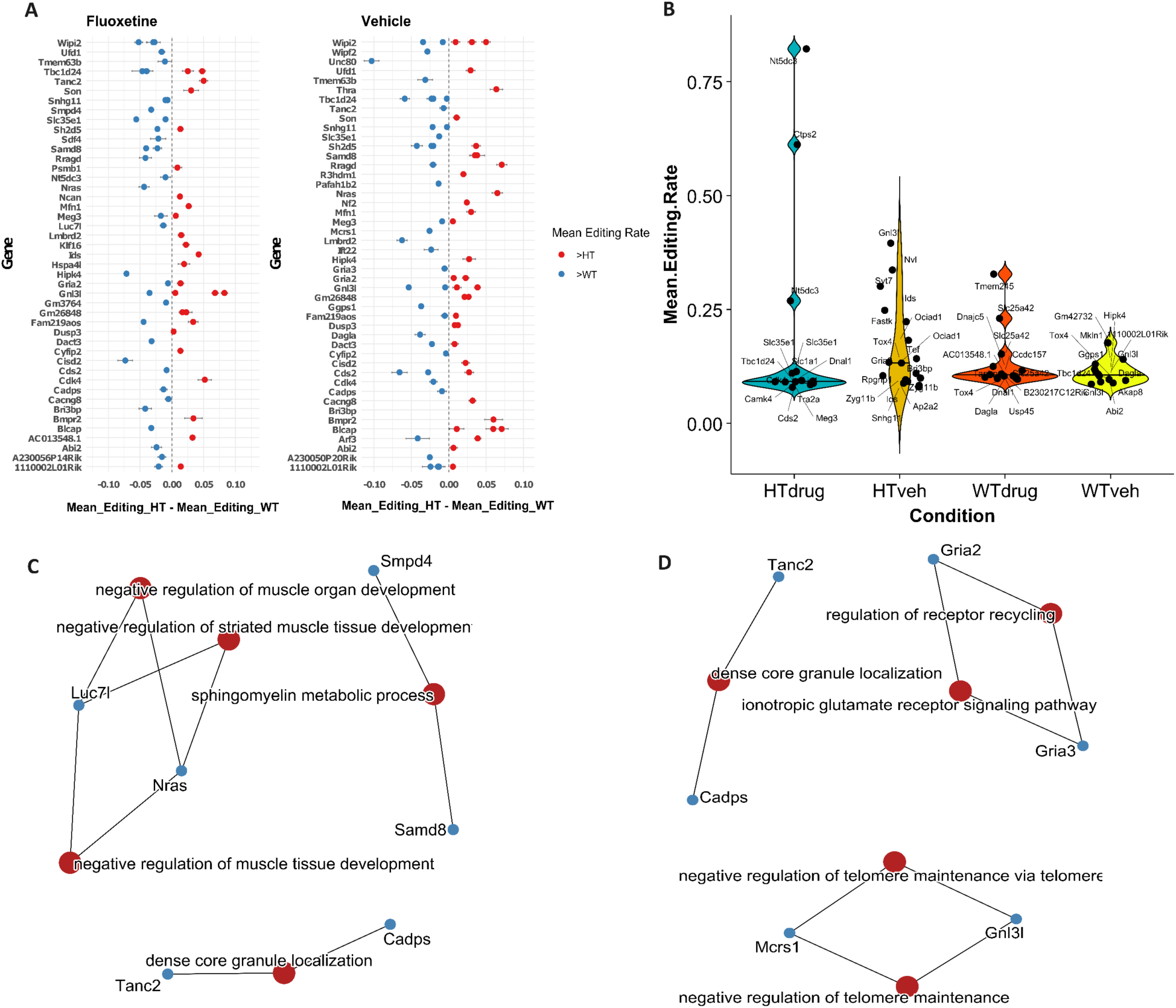
Differential ADAR Editing Between Vehicle and Fluoxetine Treated Arid1b +/- and Wild-Type Mice. **(A)** Figure shows change in editing rate for shared edited targets in the fluoxetine and vehicle treated HT, and WT mice. The shown sites were filtered for a significant 95% CI limit around the standard error (SE) for the difference between mean editing rates. The error bars shown represent upper and lower CI limits, while the x and y axis represent difference in mean editing rate for specific coordinates (chr: position), and genes mapped to those coordinates, respectively. **(B)** Genes with unique edited coordinates in all four treatment groups, where an individual coordinate is plotted against the mean editing rate in each group. Significant biological processes (BP) categories for genes with shared edited sites in **(C)** fluoxetine and **(D)** vehicle**-**treated mice, from panel **A** are shown. BP categories show that vehicle treatment promotes editing in genes related to neuronal signaling independent of the genotype.

Other than differential editing at shared targets, uniquely edited ADAR targets were also observed in each condition (Figure 2-B). We found 14 edited targets mapped to 13 genes, 17 targets mapped to 15 genes, 13 targets mapped to 11 genes, and 12 targets mapped to 11 genes in HT-drug, HT-veh, WT-drug and WT-veh groups, respectively (Supplemental Tables 4 & 5). With respect to unique edited targets, HT-drug group experienced a relative increase in site-specific editing rate, while an increase in the number of edited targets was observed in HT-veh, compared to other groups (Figure 2-B). Although the majority of the observed edited targets are overlapping (Figure 1-A), the presence of exclusively edited genes and coordinates (Figure 2-B, Supplemental Tables 4 & 5) and differential editing rates at shared coordinates (Figure 2-A) in each condition indicate nuanced plasticity in trends of ADAR editing. Interestingly, uniquely edited genes in HT-drug samples were enriched in biological processes related to neuronal transmission, highlighting potential involvement of ADAR editing in response to SSRIs (Supplemental Figure 2). For the identification of functionally important edited sites and their potential association with ASD related phenotypes, we compared our differentially edited shared and unique gene list with the SFARI gene (Abrahams et al., 2013) mouse and human database, and evolutionarily conserved (EC) ADAR targets in mammals (Pinto et al., 2014). Table 1 shows the details of the identified risk, and conserved gene targets.

**Table 1.**
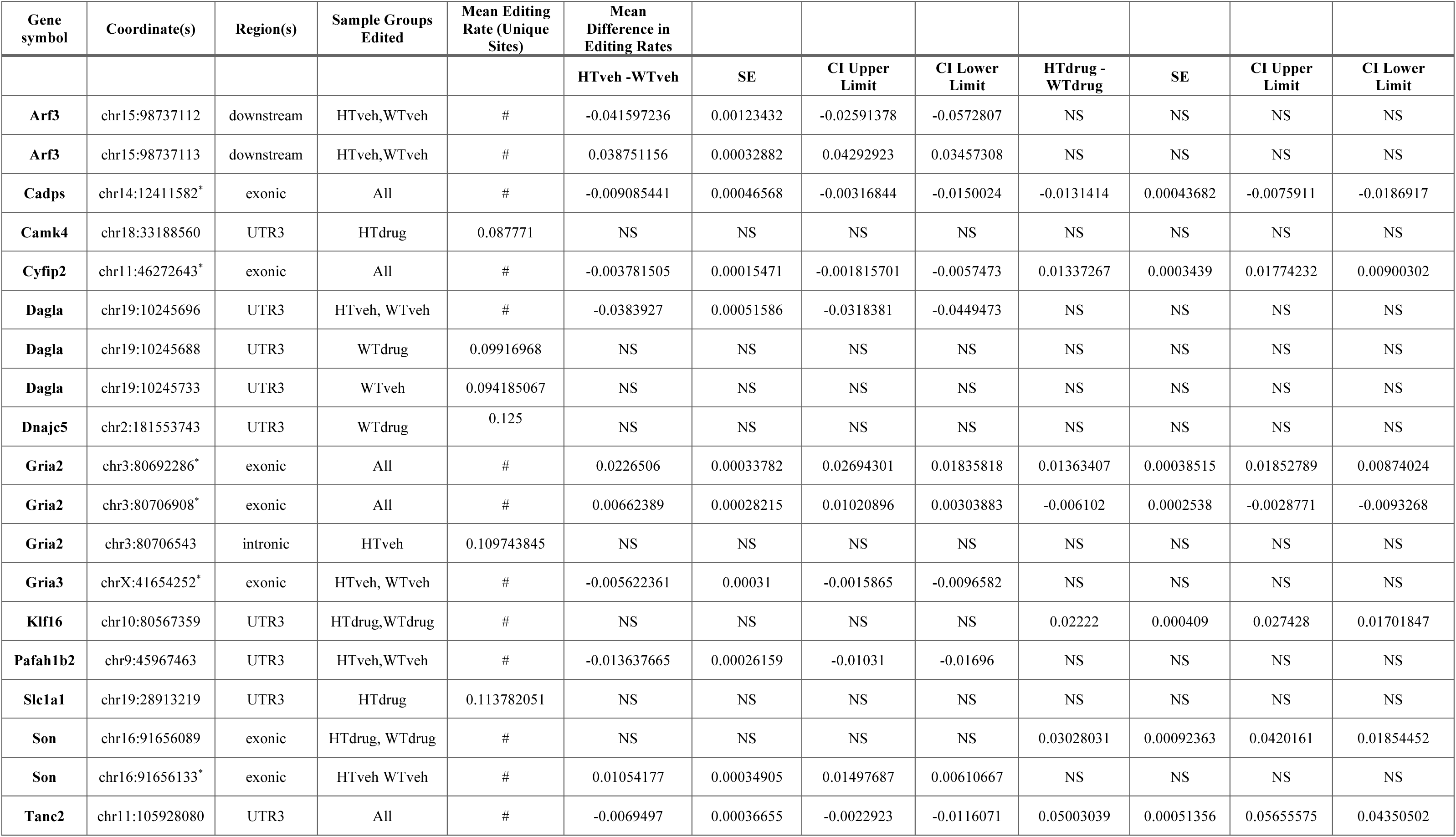
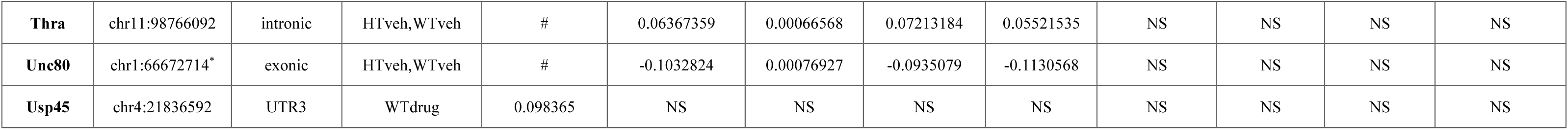
Differentially Edited ASD Associated Risk Genes. The table lists differentially edited coordinates in ASD risk genes defined by the simons foundation autism research initiative (SFARI) gene database (Abrahams et al., 2013) of human and mouse models. Columns shown display the following information: **Coordinate:** genomic coordinate with A > G or T (U) > C edit. **Region:** Region of the gene harboring the A > G or T (U) > C change. **Sample Groups Edited:** Sample groups with the specific editing changes. **Mean Difference in Editing Rate:** Increased or decreased mean editing in HT or WT mice as x-y, the “-” symbol represents increased editing in y, while “#” indicates no editing at the coordinate in the respective group. **SE:** Standard error for the difference of means. **CI Upper Limit:** Upper limit for 95% Confidence Interval around the difference of means. **CI Lower Limit:** Lower limit for 95% Confidence Interval around the difference of means. NS indicates that the coordinate is not significantly edited, while asterisk (*) indicates an evolutionarily conserved (EC) coordinate in mammals.

### ADAR Editing Alters the Number and Type of miRNAs Binding to Genes

RNA22 (v2) (Miranda et al., 2006) with default parameters was used to identify miRNA binding targets for differentially edited genes. Identified targets were then filtered for p-value ≤ 0.05, and to include the edited base position in the miRNA-mRNA hybrid. Our results show a change in the number of genes targeted by miRNAs and the number of miRNAs binding to the same gene (Supplemental Tables 6-9). For genes with shared edited sites in HT-drug and WT-drug groups, a difference in both the number of miRNA targets in edited (A > G or T(U) > C change) versus unedited states, and the number of genes being targeted, was observed (Figure 3-A, B). A comparison between the number and type of miRNA targets and the strength of binding shows that editing not only influences the type of miRNA binding to a gene target but the strength of binding for the same miRNA-mRNA hybrid as well, suggesting a role of ADAR editing in transcriptome regulation/dysregulation in ASD and response to SSRIs. Similar results were observed for the number and type of miRNA targets, type of gene target, and the strength of miRNA-mRNA binding in HT-veh, and WT-veh samples (Figure 3-C, D).

**Figure 3.**
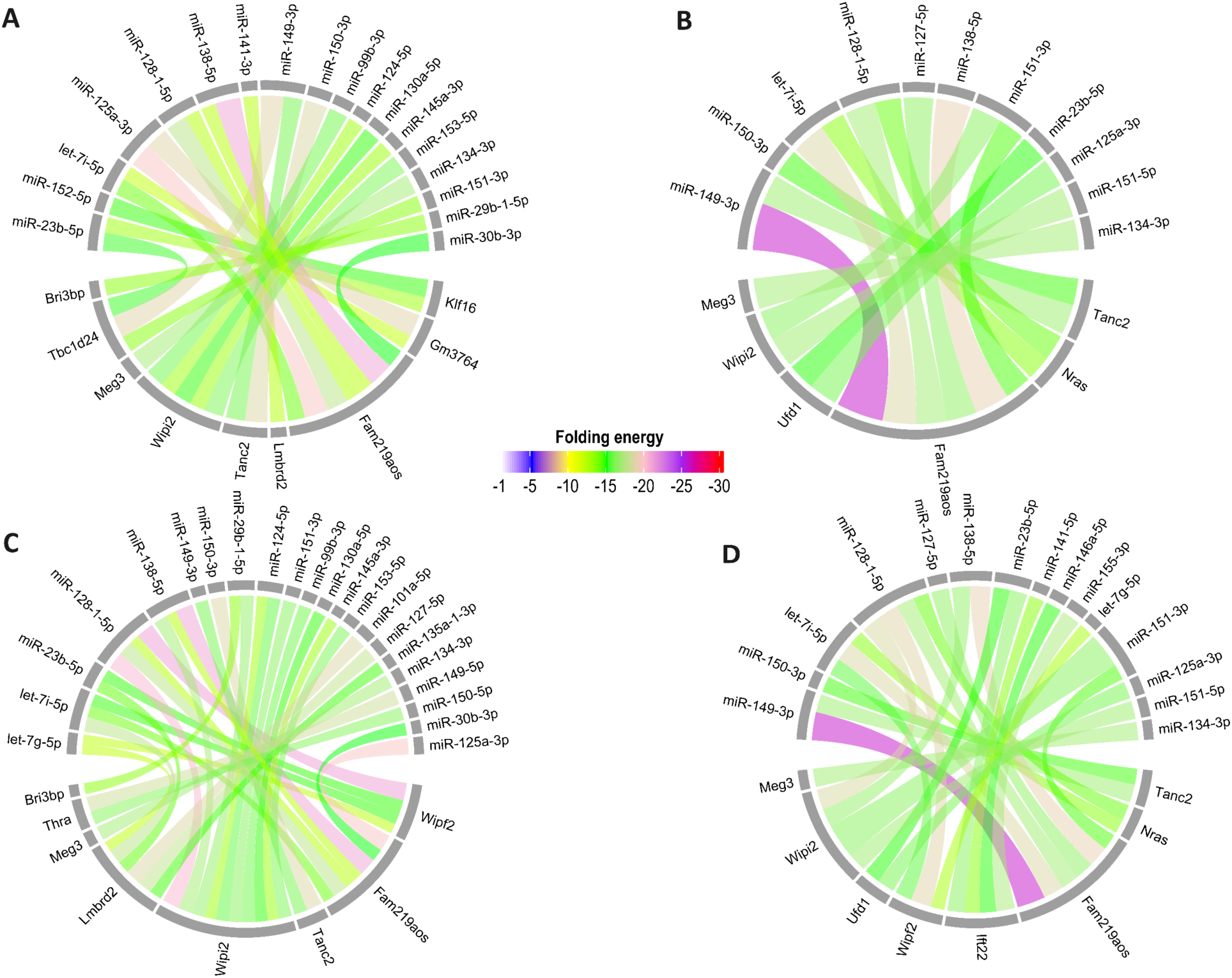
Comparison of the Number of miRNA Targets for Differentially Edited Shared Genes. Chord diagrams representing miRNA targets for genes with shared edited coordinates in **(A)** HT and WT mice treated with fluoxetine, when an A > G or T(U) > C change is present, or **(B)** absent, and **(C)** HT and WT mice treated with vehicle when an A > G or T(U) > C change is present, or **(D)** absent. The strength of the miRNA-mRNA duplex is represented by folding energy values, shown by the color key. Folding energy indicates the thermodynamic stability of the miRNA-mRNA duplex, where lower values indicate a more stable interaction (see methods for details).

For instance, Meg3 is a miRNA target in both edited and unedited states, but the type of miRNA binding to the gene is different when edited (Figure 3-C, D). Similarly, for Fam219aos, identical miRNAs are binding to the gene in edited and unedited states, but the difference in folding energy values (e.g., for let-7i-5p) indicates a change in binding strength (Figure 3-C, D). Some genes, such as Klf16 and Bri3bp, were targeted by miRNAs only when edited, while the Nras gene showed miRNA binding only when unedited (Figure 3). Next, we compared miRNA targets for uniquely edited genes in HT, and WT mice. Given the smaller number of unique edited sites, only a few significant miRNA-mRNA interactions were observed (Supplemental Tables 8 & 9).

### Differential Gene Expression Provides a Limited Overview of the Transcriptome Dysregulation

To investigate whether editing dysregulation relates to differential gene expression, we compared our lists of DEGs and differentially edited genes (Supplemental Tables 1B,1C & 11) and found no overlap. We further performed GO analyses for the DEGs, and edited gene lists with significant miRNA binding targets. As shown in Figure 4 (B, C), GO terms show a drastically different ontology for the compared gene lists. These findings highlight the limitation of differential gene expression analysis alone to identify dysregulations in the transcriptome. None of the ADAR genes (Adar1, Adar2, or Adar3) was found differentially edited according to our set thresholds. However, to evaluate precise differences between ADAR gene counts that may explain the differential ADAR editing landscape of the compared groups, normalized transcript count values for all ADAR genes were compared (Figure 4-A). It should be noted that ADAR editing changes are dynamic and can depend on other factors besides ADAR expression, such as interaction of ADARs with dsRNA-binding proteins (Deffit and Hundley, 2016; Jacobs et al., 2009; Tan et al., 2017). For a direct comparison, the fluoxetine-treated HT mice were only compared with fluoxetine-treated WT mice, while vehicle-treated HT mice were only compared with vehicle-treated WT mice. Our results show no significant difference in ADAR expression except Adar1 (Figure 4-A), where Adar1 is upregulated in HT-drug (padj-value = 0.01). However, no significant difference between Adar1 isoforms (Adarp150, Adarp110) was found (Supplemental Figure 1-B, C).

**Figure 4.**
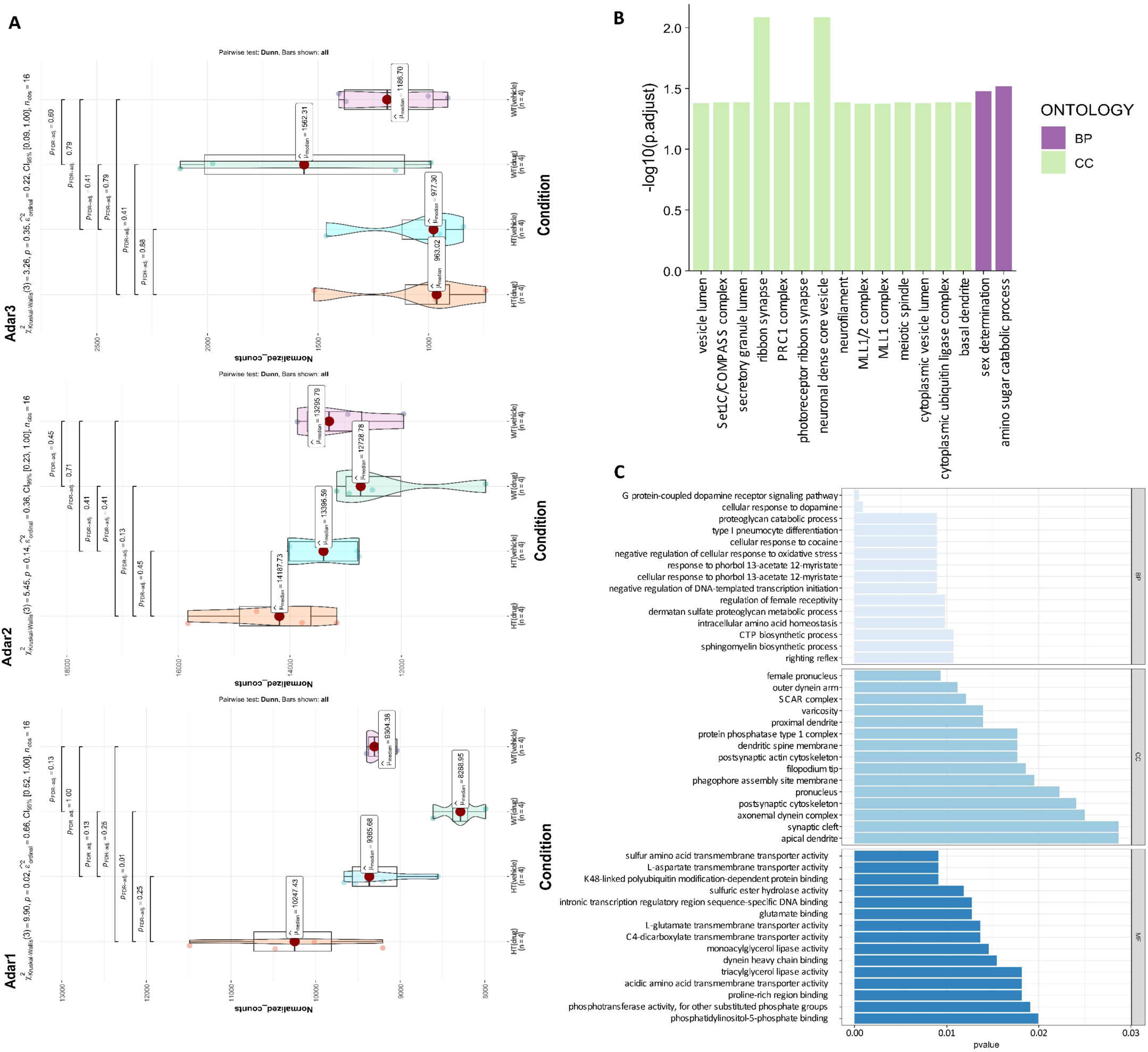
Comparison of Transcriptome Dysregulation in Terms of Differential Gene Expression and ADAR Editing. **(A)** Comparison of gene counts for Adar1, Adar2, and Adar3 between HT and WT mice treated with fluoxetine or vehicle. **(B)** Gene Ontology (GO) terms for differentially expressed genes (DEGs) between HT and WT mice treated with fluoxetine or vehicle. Color key represents significant GO terms, Biological Process (BP), and Cellular Component (CC). No significant Molecular Function (MF) categories were found for DEGs. **(C)** Gene Ontology (GO) terms for genes with differential editing and miRNA binding targets between HT and WT mice treated with fluoxetine or vehicle. Significant GO terms are shown as Biological Process (BP), Molecular Function (MF), and Cellular Component (CC).

## Discussion

Phenotypic heterogeneity among autistic individuals highlights a complex, multifactorial genetic and environmental influence on the disease manifestation (Geschwind, 2009; Lord et al., 2020). A previous study has reported differential ADAR editing in human postmortem brain samples from autistic individuals, where a trend towards a decrease in editing rate was observed (Tran et al., 2019). Here we present findings that differential ADAR activity is associated with ASD and use of SSRIs, in terms of change in the number of edited sites, rate of editing per site, and the effect of editing on miRNA binding to gene targets. While further experimental studies are needed to fully understand this phenomenon and its implications for development of ASD diagnostics and treatment, our study provides an initial subset of editing signatures that can be tested in future studies.

Using the set of experimentally validated ADAR-edited sites from the REDIportal (Picardi et al., 2017) as a reference, we demonstrate that use of SSRIs is associated with transcriptome-wide differential ADAR editing changes in both autistic and wild-type phenotypes. Although not statistically significant, we found a global trend for an increase in editing in terms of the number of edited targets and site-specific editing rate in HT-veh mice compared to other groups (Supplemental Figure1). These differences from Tran and colleagues (Tran et al., 2019) could potentially arise from inherent differences in the mouse and human genomes (Yue et al., 2014), or the use of specific autistic mouse models (Provenzano et al., 2016). Kim and colleagues (Kim et al., 2022) have reported behavioral improvements closer to wild-type in autistic mice after chronic fluoxetine treatment. In our analysis, fluoxetine treatment was shown to increase and decrease the number of edited sites in WT and HT mice, respectively (Supplemental Figure1), suggesting a potential role of ADAR editing in mediating the therapeutic effects of fluoxetine. Whereas changes in site-specific editing rate (Figure 2-E) suggest a more nuanced, phenotype independent effect of fluoxetine on editing rate. Fluoxetine treatment has been shown to increase Adar2 expression in cell cultures and intact mouse brain with no difference in Adar1 expression (Li et al., 2011). In our analysis, no ADAR gene was differentially expressed except for Adar1, where DEseq2 normalized count values were relatively higher in HT-drug mice (Figure 4-A). These differences could arise due to the type of gene expression method used in the (Li et al., 2011) study, the type of cells and/or tissues, and the dose of fluoxetine (Tate et al., 2021). Moreover, higher expression of Adar1 may explain the comparatively distinct editing landscape of HT-drug mice (Figure 1-E).

It is important to note that ADAR editing predominantly occurs in non-coding regions, and recoding-type editing represents only a small, albeit important fraction of editing activity (Gabay et al., 2022; Nishikura, 2016; Yang et al., 2013). In line with that, we found the majority of differences in editing activity between HT and WT mice located in non-coding regulatory regions such as 3’ and 5’UTRs, ncRNA, and introns. To identify functionally important sites in coding regions, we used a set of known EC editing targets in mammals (Pinto et al., 2014). When compared with conserved recoding sites, we found 12 targets edited in vehicle-treated HT and WT mice, while 8 of those targets were edited in fluoxetine treated HT and WT mice (Supplemental Table 10), including two sites in the Gria2, also commonly known as GluA2, gene. Gria2 encodes the GluA2 subunit of α-amino-3-hydroxy-5-methyl-4-isoxazolepropionic acid receptor (AMPAR) ionotropic glutamate receptors, which regulate excitatory synapses (Isaac et al., 2007). ADAR editing at the Q607R site in Gria2 mRNA regulates the excitatory synapse by rendering the channel impermeable to calcium (Ca^2+^) (Isaac et al., 2007).

Gria2 is also an ASD associated risk gene (Cai et al., 2022), and variants of Gria2 and alterations in RNA editing have been linked with other neurological phenotypes such as embryonic lethality (Higuchi et al., 2000), and Alzheimer’s disease (Wright et al., 2023). The therapeutic effects of fluoxetine hydrochloride are mediated through cellular events that promote neurogenesis and synaptic plasticity (Ampuero et al., 2013, 2010; Wang et al., 2011). AMPAR receptors, including GluA2, can modify the strength of synapses during bidirectional synaptic (excitation or depression) plasticity events (Diering and Huganir, 2018). A study in rats has shown that fluoxetine treatment can increase the expression of Gria2 in the hippocampus (Ampuero et al., 2013). Although not differentially expressed, Gria2 was edited in all conditions, including two EC sites (Supplemental Table 10) and a unique edited site (Supplemental Table 4), in HT-veh mice. These findings suggest the use of SSRI, potentially dose-dependent (Tate et al., 2021), can affect transcriptome regulation in multiple ways. It is noteworthy that one EC site in Gria2 (chr3: 80706908) resides in exon 11, which is the same exon in the mm10 genome assembly, harboring the Q/R site (Perez et al., 2024). Feldmeyer and colleagues (Feldmeyer et al., 1999) have reported that mutant mice with differential editing levels at the Gria2 Q/R site and differences in expression of Gria2 mRNA showed diverse phenotypes ranging from epileptic seizures to premature death. This indicates functional consequences of not only Q/R editing, but also the availability of Gria2 mRNA in the cell. Moreover, ADAR editing can influence mRNA stability and expression through interactions with other RNA-binding proteins (Wang et al., 2013), suggesting potential consequences of Gria2 mRNA editing other than the Q/R site. Although we found no significant change in Q/R (chr3: 80706912) site editing, in one of the WT-drug (SRR19308799) samples (Supplemental Table 13) editing at Q/R sites was ∼ 98% compared to ∼99% in other samples. While seemingly minor, this under editing difference (assuming its magnitude is correctly reflected in the transcriptome data) could result in an increase in the number of AMPA receptors being permeable to Ca^2+^, in turn leading to excitotoxicity, and subsequent neuronal and synaptic dysfunction (Konen et al., 2020; Wright et al., 2023). This highlights the need to further investigate the effects of SSRI use on Gria2 editing with broader sampling of tissues and dose ranges.

Another ASD associated risk gene, Camk4 (Abrahams et al., 2013), was uniquely edited in the HT-drug group (Supplemental Table 4). Following antidepressant treatment, including fluoxetine, Camk4, or calcium/calmodulin-dependent kinase IV, has been shown to promote neurogenesis by increasing the phosphorylation of transcription factor cAMP-response element-binding protein (CREB) (Song et al., 2013; Tiraboschi et al., 2004). Although the effect of nucleotide changes by ADARs on RNA stability and expression is less understood, the ADAR function is shown to influence alternative splicing, binding of dsRNA binding proteins, and secondary structural changes in the mRNA (Solomon et al., 2017, 2013; Wang et al., 2013). Moreover, a recent study has shown that base changes in untranslated regions and coding sequence of mRNA could affect RNA secondary structure formation and hence mRNA expression (Mauger et al., 2019). Based on the presented evidence, differential editing could potentially influence post-transcriptional regulation of mRNA in terms of lowering gene expression, and changes in mRNA secondary structure.

Unique edited sites in HT-drug and WT-drug (Supplemental Tables 4 & 5) indicate phenotype-specific effects of fluoxetine treatment. We found another EC site in the Dact3 transcript that was differentially edited, with an increase in editing in WT mice for both fluoxetine and vehicle treatments (Figure 2-A, Supplemental Table 10). Dact3 expression is essentially important during early neurodevelopment (Fisher et al., 2006), and an overall increase in Dact3 editing at this conserved site has been reported in fetal brain after poly I:C induction in MIA models for neurodevelopmental disease (Tsivion-Visbord et al., 2020). Compared to these findings, the observed relative increase in Dact3 editing at the conserved sites in both WT-drug and WT-veh mice in our analysis indicates complex dynamics of ADAR editing in response to environmental stimuli, individual phenotypes, and developmental stage. For another conserved site in the Blcap gene, fluoxetine treatment caused a decrease in editing in HT mice, while editing in WT mice was increased (Supplemental Table 10). As shown in Figure 1-C, one shared exonic edited site between HT-drug and HT-veh maps to the Mfn1 (chr3:32561485) gene, with a relative decrease in editing in HT-drug (Supplemental Table 14). These findings provide additional evidence and targets for the effects of SSRIs beyond neuronal processes (Wang et al., 2026; Yuan et al., 2023).

Two more genes, Nt5dc3 and Sdf4, with differentially edited sites in only the fluoxetine-treated groups, had decreased editing in HT as compared to WT mice at the same site (Figure 2-A). Nt5dc3 is a transmembrane protein with largely uncharacterized function (Cros-Perrial and Jordheim, 2025). Differential expression of Nt5dc3 has been shown in reversal learning behavior in mice, suggesting its role in impulse control and behavioral disorders (Laughlin et al., 2011). Sdf4 encodes a calcium-binding signaling protein, and alternative splicing of the mRNA produces three protein isoforms with distinct cellular localizations and functions (Honoré, 2009; Zhu et al., 2021). Both Nt5dc3 and Sdf4 have differential expression in schizophrenia and bipolar disorder (Zhang et al., 2020). This points towards a complex yet overlapping transcriptome regulation in diverse behavioral phenotypes and highlights the need to study cellular and physiological impacts of transcriptome modifications such as ADAR editing in complex phenotypes and disorders.

Through RNA editing, two versions (edited and unedited) of the same RNA molecule can be preserved and translated at the given time (Goldeck et al., 2022). This not only contributes to transcriptome diversity, but can also impact mRNAs interaction with other transcriptional regulatory elements such as miRNAs (Park et al., 2021; Roberts et al., 2018; Tomaselli et al., 2013). We observed differences in miRNA binding to genes in their edited versus unedited state. Notably, Klf16 and Thra, some of the ASD associated risk genes (Abrahams et al., 2013), were targeted by miRNAs only when edited. Klf16 is a transcriptional repressor and is highly expressed in the developing brain (McConnell and Yang, 2010; Syafruddin et al., 2020). In our analysis, Klf16 has a shared edited site in its 3’UTR only in HT-drug and WT-drug (Figure 2-A), and is targeted by multiple miRNAs when edited (Figure 3-A). Differentially edited genes targeted by miRNAs were significantly enriched in biological processes related to transcriptional regulation and neurotransmission (Figure 4-C). Another study has reported that gene targets for highly expressed miRNA in the brain in ASD are also involved in the aforementioned related categories (Wong et al., 2022). In our analysis, none of the edited genes with differential miRNA binding had differential expression, including ASD associated risk genes, and were enriched for different GO terms (Figure 4). These findings highlight the narrow scope of differential gene expression analysis, and underscore the need to consider other layers of transcriptome regulation, including ADAR editing, in transcriptome studies and therapeutic clinical trials. In conclusion, our study identifies a subset of potential ADAR targets associated with the use of SSRIs. Although our computational analyses cannot determine the definitive causal relationships, nonetheless, the presence of unique edited sites and differentially edited sites in known ASD risk genes that are impacted by the SSRIs use can be interpreted as indicators of their potential contribution to mechanisms underlying disease pathology. Additionally, the differentially edited sites overlapped with conserved ADAR targets in mammals, suggesting important functional implications of dynamic editing changes. In addition to adding or eliminating miRNA targets, differences in the number, type, and strength of miRNA binding to a gene, in the edited and unedited state, further support a potential role of editing in mRNA stability (Bagga et al., 2005; Valinezhad Orang et al., 2014). Moreover, the enrichment of differentially edited genes in biological processes related to neurotransmission indicates that autism related phenotypes span beyond social behavior features and further emphasizes the role of ADAR editing in neurodevelopmental disorders. This study is limited by the lack of experimental validation and a sex bias, because only male animals were used in the original dataset (Kim et al., 2022). Future studies should focus on the experimental validation of these variants of “uncertain significance” (Richards et al., 2015) to elucidate their role in the pathophysiology of complex neurodevelopmental disorders, and expand the experiments to include both male and female animals.

## Acknowledgement

We would like to thank Dr. Kim Woo Yang, PhD, and members of the Piontkivska lab for their assistance with this study.

## Conflict of Interest

The authors declare no conflict of interest.

## Data availability

The dataset used in this study is publicly available in the NCBI SRA/BioProject repository, as BioProject PRJNA839604 (Kim et al., 2022), https://www.ncbi.nlm.nih.gov/bioproject/PRJNA839604. Supplementary Figures and Tables, and R code used in the analysis can be found at https://github.com/RNAdetective/Site-Specific_ADAR_Editing_in_Arid1b_mouse.

## List of abbreviations

ADAR: Adenosine Deaminase Acting on RNA
ASD: Autism Spectrum Disorder
AIDD: Automated Isoform Diversity Detector
BP: Base Pair
DEGs: Differentially Expressed Genes
dsRNA: Double-Stranded RNA
HT: Heterozygous
ISG: Interferon Stimulated Gene
miRNA: Micro RNA
MIA: Maternal Immune Activation
ncRNA: Non-Coding RNA
PFC: Prefrontal Cortex
Poly I:C: Polyinosinic Polycytidylic Acid
SERT: Serotonin Transporter
SSRIs: Selective Serotonin Reuptake Inhibitors
WT: Wild Type

## Notes

https://github.com/RNAdetective/Site-Specific_ADAR_Editing_in_Arid1b_mouse

### Competing Interest Statement

The authors have declared no competing interest.

## References

Abrahams, B.S., Arking, D.E., Campbell, D.B., Mefford, H.C., Morrow, E.M., Weiss, L.A., Menashe, I., Wadkins, T., Banerjee-Basu, S., Packer, A., 2013. SFARI Gene 2.0: a community-driven knowledgebase for the autism spectrum disorders (ASDs). Mol. Autism 4, 36. 10.1186/2040-2392-4-36

Ampuero, E., Rubio, F.J., Falcon, R., Sandoval, M., Diaz-Veliz, G., Gonzalez, R.E., Earle, N., Dagnino-Subiabre, A., Aboitiz, F., Orrego, F., Wyneken, U., 2010. Chronic fluoxetine treatment induces structural plasticity and selective changes in glutamate receptor subunits in the rat cerebral cortex. Neuroscience 169, 98–108. 10.1016/j.neuroscience.2010.04.035

Ampuero, E., Stehberg, J., Gonzalez, D., Besser, N., Ferrero, M., Diaz-Veliz, G., Wyneken, U., Rubio, F.J., 2013. Repetitive fluoxetine treatment affects long-term memories but not learning. Behav. Brain Res. 247, 92–100. 10.1016/j.bbr.2013.03.011

Atladóttir, H.Ó., Thorsen, P., Østergaard, L., Schendel, D.E., Lemcke, S., Abdallah, M., Parner, E.T., 2010. Maternal Infection Requiring Hospitalization During Pregnancy and Autism Spectrum Disorders. J. Autism Dev. Disord. 40, 1423–1430. 10.1007/s10803-010-1006-y

Bagga, S., Bracht, J., Hunter, S., Massirer, K., Holtz, J., Eachus, R., Pasquinelli, A.E., 2005. Regulation by let-7 and lin-4 miRNAs Results in Target mRNA Degradation. Cell 122, 553–563. 10.1016/j.cell.2005.07.031

Bass, B.L., 2002. RNA Editing by Adenosine Deaminases That Act on RNA. Annu. Rev. Biochem. 71, 817–846. 10.1146/annurev.biochem.71.110601.135501

Behm, M., Öhman, M., 2016. RNA Editing: A Contributor to Neuronal Dynamics in the Mammalian Brain. Trends Genet. 32, 165–175. 10.1016/j.tig.2015.12.005

Burns, C.M., Chu, H., Rueter, S.M., Hutchinson, L.K., Canton, H., Sanders-Bush, E., Emeson, R.B., 1997. Regulation of serotonin-2C receptor G-protein coupling by RNA editing. Nature 387, 303–308. 10.1038/387303a0

Cai, Q., Zhou, Z., Luo, R., Yu, T., Li, D., Yang, F., Yang, Z., 2022. Novel GRIA2 variant in a patient with atypical autism spectrum disorder and psychiatric symptoms: a case report. BMC Pediatr. 22, 629. 10.1186/s12887-022-03702-7

Cha, J.H., Kinsman, S.L., Johnston, M.V., 1994. RNA editing of a human glutamate receptor subunit. Brain Res. Mol. Brain Res. 22, 323–328. 10.1016/0169-328x(94)90061-2

Choi, L., An, J.-Y., 2021. Genetic architecture of autism spectrum disorder: Lessons from large-scale genomic studies. Neurosci. Biobehav. Rev. 128, 244–257. 10.1016/j.neubiorev.2021.06.028

Cros-Perrial, E., Jordheim, L.P., 2025. A phenotypic journey into NT5DC proteins in cancer and other diseases. Exp. Cell Res. 446, 114468. 10.1016/j.yexcr.2025.114468

Deffit, S.N., Hundley, H.A., 2016. To edit or not to edit: regulation of ADAR editing specificity and efficiency. WIREs RNA 7, 113–127. 10.1002/wrna.1319

Diering, G.H., Huganir, R.L., 2018. The AMPA receptor code of synaptic plasticity. Neuron 100, 314–329. 10.1016/j.neuron.2018.10.018

Dracheva, S., Patel, N., Woo, D.A., Marcus, S.M., Siever, L.J., Haroutunian, V., 2008. Increased serotonin 2C receptor mRNA editing: a possible risk factor for suicide. Mol. Psychiatry 13, 1001–1010. 10.1038/sj.mp.4002081

Eissa, N., Sadeq, A., Sasse, A., Sadek, B., 2020. Role of Neuroinflammation in Autism Spectrum Disorder and the Emergence of Brain Histaminergic System. Lessons Also for BPSD? Front. Pharmacol. 11, 886. 10.3389/fphar.2020.00886

Englander, M.T., Dulawa, S.C., Bhansali, P., Schmauss, C., 2005. How Stress and Fluoxetine Modulate Serotonin 2C Receptor Pre-mRNA Editing. J. Neurosci. 25, 648–651. 10.1523/JNEUROSCI.3895-04.2005

Eran, A., Li, J.B., Vatalaro, K., McCarthy, J., Rahimov, F., Collins, C., Markianos, K., Margulies, D.M., Brown, E.N., Calvo, S.E., Kohane, I.S., Kunkel, L.M., 2013. Comparative RNA editing in autistic and neurotypical cerebella. Mol. Psychiatry 18, 1041–1048. 10.1038/mp.2012.118

Feldmeyer, D., Kask, K., Brusa, R., Kornau, H.-C., Kolhekar, R., Rozov, A., Burnashev, N., Jensen, V., Hvalby, Ø., Sprengel, R., Seeburg, P.H., 1999. Neurological dysfunctions in mice expressing different levels of the Q/R site–unedited AMPAR subunit GluR–B. Nat. Neurosci. 2, 57–64. 10.1038/4561

Fisher, D.A., Kivimäe, S., Hoshino, J., Suriben, R., Martin, P., Baxter, N., Cheyette, B.N.R., 2006. Three Dact gene family members are expressed during embryonic development and in the adult brains of mice. Dev. Dyn. 235, 2620–2630. 10.1002/dvdy.20917

Fuller, R.W., Wong, D.T., Robertson, D.W., 1991. Fluoxetine, a selective inhibitor of serotonin uptake. Med. Res. Rev. 11, 17–34. 10.1002/med.2610110103

Gabay, O., Shoshan, Y., Kopel, E., Ben-Zvi, U., Mann, T.D., Bressler, N., Cohen-Fultheim, R., Schaffer, A.A., Roth, S.H., Tzur, Z., Levanon, E.Y., Eisenberg, E., 2022. Landscape of adenosine-to-inosine RNA recoding across human tissues. Nat. Commun. 13, 1184. 10.1038/s41467-022-28841-4

Gallo, A., Vukic, D., Michalík, D., O’Connell, M.A., Keegan, L.P., 2017. ADAR RNA editing in human disease; more to it than meets the I. Hum. Genet. 136, 1265–1278. 10.1007/s00439-017-1837-0

Garrido-Torres, N., Guzmán-Torres, K., García-Cerro, S., Pinilla Bermúdez, G., Cruz-Baquero, C., Ochoa, H., García-González, D., Canal-Rivero, M., Crespo-Facorro, B., Ruiz-Veguilla, M., 2024. miRNAs as biomarkers of autism spectrum disorder: a systematic review and meta-analysis. Eur. Child Adolesc. Psychiatry 33, 2957–2990. 10.1007/s00787-023-02138-3

GENCODE - Mouse Release M10 [WWW Document], 2024. URL https://www.gencodegenes.org/mouse/release_M10.html (accessed 2.6.25).

Genovese, A., Butler, M.G., 2023. The Autism Spectrum: Behavioral, Psychiatric and Genetic Associations. Genes 14, 677. 10.3390/genes14030677

Geschwind, D.H., 2009. Advances in Autism. Annu. Rev. Med. 60, 367–380. 10.1146/annurev.med.60.053107.121225

Ghoshal, A., Shankar, R., Bagchi, S., Grama, A., Chaterji, S., 2015. MicroRNA target prediction using thermodynamic and sequence curves. BMC Genomics 16, 999. 10.1186/s12864-015-1933-2

Goldeck, M., Gopal, A., Jantsch, M.F., Mansouri Khosravi, H.R., Rajendra, V., Vesely, C., 2022. How RNA editing keeps an I on physiology. Am. J. Physiol.-Cell Physiol. 323, C1496– C1511. 10.1152/ajpcell.00191.2022

Griffiths-Jones, S., Grocock, R.J., van Dongen, S., Bateman, A., Enright, A.J., 2006. miRBase: microRNA sequences, targets and gene nomenclature. Nucleic Acids Res. 34, D140– D144. 10.1093/nar/gkj112

Gurevich, I., Tamir, H., Arango, V., Dwork, A.J., Mann, J.J., Schmauss, C., 2002. Altered editing of serotonin 2C receptor pre-mRNA in the prefrontal cortex of depressed suicide victims. Neuron 34, 349–356. 10.1016/s0896-6273(02)00660-8

Gzieło, K., Piotrowska, D., Litwa, E., Popik, P., Nikiforuk, A., 2023. Maternal immune activation affects socio-communicative behavior in adult rats. Sci. Rep. 13, 1918. 10.1038/s41598-023-28919-z

Haddad, F.L., Patel, S.V., Schmid, S., 2020. Maternal Immune Activation by Poly I:C as a preclinical Model for Neurodevelopmental Disorders: A focus on Autism and Schizophrenia. Neurosci. Biobehav. Rev. 113, 546–567. 10.1016/j.neubiorev.2020.04.012

Han, V.X., Patel, S., Jones, H.F., Dale, R.C., 2021. Maternal immune activation and neuroinflammation in human neurodevelopmental disorders. Nat. Rev. Neurol. 17, 564–579. 10.1038/s41582-021-00530-8

Hazra, A., 2017. Using the confidence interval confidently. J. Thorac. Dis. 9, 4125–4130. 10.21037/jtd.2017.09.14

Higuchi, M., Maas, S., Single, F.N., Hartner, J., Rozov, A., Burnashev, N., Feldmeyer, D., Sprengel, R., Seeburg, P.H., 2000. Point mutation in an AMPA receptor gene rescues lethality in mice deficient in the RNA-editing enzyme ADAR2. Nature 406, 78–81. 10.1038/35017558

Honoré, B., 2009. The rapidly expanding CREC protein family: members, localization, function, and role in disease. BioEssays 31, 262–277. 10.1002/bies.200800186

Horackova, H., Karahoda, R., Cerveny, L., Vachalova, V., Ebner, R., Abad, C., Staud, F., 2021. Effect of Selected Antidepressants on Placental Homeostasis of Serotonin: Maternal and Fetal Perspectives. Pharmaceutics 13, 1306. 10.3390/pharmaceutics13081306

Huntley, M.A., Lou, M., Goldstein, L.D., Lawrence, M., Dijkgraaf, G.J.P., Kaminker, J.S., Gentleman, R., 2016. Complex regulation of ADAR-mediated RNA-editing across tissues. BMC Genomics 17, 61. 10.1186/s12864-015-2291-9

Hwang, T., Park, C.-K., Leung, A.K.L., Gao, Y., Hyde, T.M., Kleinman, J.E., Rajpurohit, A., Tao, R., Shin, J.H., Weinberger, D.R., 2016. Dynamic regulation of RNA editing in human brain development and disease. Nat. Neurosci. 19, 1093–1099. 10.1038/nn.4337

Isaac, J.T.R., Ashby, M.C., McBain, C.J., 2007. The Role of the GluR2 Subunit in AMPA Receptor Function and Synaptic Plasticity. Neuron 54, 859–871. 10.1016/j.neuron.2007.06.001

Iwamoto, K., Kato, T., 2003. RNA editing of serotonin 2C receptor in human postmortem brains of major mental disorders. Neurosci. Lett. 346, 169–172. 10.1016/s0304-3940(03)00608-6

Jacobs, M.M., Fogg, R.L., Emeson, R.B., Stanwood, G.D., 2009. ADAR1 and ADAR2 Expression and Editing Activity during Forebrain Development. Dev. Neurosci. 31, 223–237. 10.1159/000210185

Jung, E.-M., Moffat, J.J., Liu, J., Dravid, S.M., Gurumurthy, C.B., Kim, W.-Y., 2017. Arid1b haploinsufficiency disrupts cortical interneuron development and mouse behavior. Nat. Neurosci. 20, 1694–1707. 10.1038/s41593-017-0013-0

Kim, Hyosang, Kim, D., Cho, Y., Kim, K., Roh, J.D., Kim, Y., Yang, E., Kim, S.S., Ahn, S., Kim, Hyun, Kang, H., Bae, Y., Kim, E., 2022. Early postnatal serotonin modulation prevents adult-stage deficits in Arid1b-deficient mice through synaptic transcriptional reprogramming. Nat. Commun. 13, 5051. 10.1038/s41467-022-32748-5

Konen, L.M., Wright, A.L., Royle, G.A., Morris, G.P., Lau, B.K., Seow, P.W., Zinn, R., Milham, L.T., Vaughan, C.W., Vissel, B., 2020. A new mouse line with reduced GluA2 Q/R site RNA editing exhibits loss of dendritic spines, hippocampal CA1-neuron loss, learning and memory impairments and NMDA receptor-independent seizure vulnerability. Mol. Brain 13, 27. 10.1186/s13041-020-0545-1

Lan, X.-Y., Gu, Y.-Y., Li, M.-J., Song, T.-J., Zhai, F.-J., Zhang, Y., Zhan, J.-S., Böckers, T.M., Yue, X.-N., Wang, J.-N., Yuan, S., Jin, M.-Y., Xie, Y.-F., Dang, W.-W., Hong, H.-H., Guo, Z.-R., Wang, X.-W., Zhang, R., 2023. Poly(I:C)-induced maternal immune activation causes elevated self-grooming in male rat offspring: Involvement of abnormal postpartum static nursing in dam. Front. Cell Dev. Biol. 11. 10.3389/fcell.2023.1054381

Laughlin, R.E., Grant, T.L., Williams, R.W., Jentsch, J.D., 2011. Genetic Dissection of Behavioral Flexibility: Reversal Learning in Mice. Biol. Psychiatry, Nucleus Accumbens Neuroadaptations and Relapse in Addiction 69, 1109–1116. 10.1016/j.biopsych.2011.01.014

Lee, A., Choo, H., Jeon, B., 2022. Serotonin Receptors as Therapeutic Targets for Autism Spectrum Disorder Treatment. Int. J. Mol. Sci. 23, 6515. 10.3390/ijms23126515

Li, B., Zhang, S., Zhang, H., Hertz, L., Peng, L., 2011. Fluoxetine affects GluK2 editing, glutamate-evoked Ca2+ influx and extracellular signal-regulated kinase phosphorylation in mouse astrocytes. J. Psychiatry Neurosci. JPN 36, 322–338. 10.1503/jpn.100094

Li, J., Xu, X., Liu, J., Zhang, S., Tan, X., Li, Z., Zhang, J., Wang, Z., 2022. Decoding microRNAs in autism spectrum disorder. Mol. Ther. - Nucleic Acids 30, 535–546. 10.1016/j.omtn.2022.11.005

Lord, C., Brugha, T.S., Charman, T., Cusack, J., Dumas, G., Frazier, T., Jones, E.J.H., Jones, R.M., Pickles, A., State, M.W., Taylor, J.L., Veenstra-VanderWeele, J., 2020. Autism spectrum disorder. Nat. Rev. Dis. Primer 6, 5. 10.1038/s41572-019-0138-4

Love, M.I., Huber, W., Anders, S., 2014. Moderated estimation of fold change and dispersion for RNA-seq data with DESeq2. Genome Biol. 15, 550. 10.1186/s13059-014-0550-8

Maenner, M.J., 2023. Prevalence and Characteristics of Autism Spectrum Disorder Among Children Aged 8 Years — Autism and Developmental Disabilities Monitoring Network, 11 Sites, United States, 2020. MMWR Surveill. Summ. 72. 10.15585/mmwr.ss7202a1

Man, K.K.C., Tong, H.H.Y., Wong, L.Y.L., Chan, E.W., Simonoff, E., Wong, I.C.K., 2015. Exposure to selective serotonin reuptake inhibitors during pregnancy and risk of autism spectrum disorder in children: A systematic review and meta-analysis of observational studies. Neurosci. Biobehav. Rev. 49, 82–89. 10.1016/j.neubiorev.2014.11.020

Mansi, L., Tangaro, M.A., Lo Giudice, C., Flati, T., Kopel, E., Schaffer, A.A., Castrignanò, T., Chillemi, G., Pesole, G., Picardi, E., 2021. REDIportal: millions of novel A-to-I RNA editing events from thousands of RNAseq experiments. Nucleic Acids Res. 49, D1012–D1019. 10.1093/nar/gkaa916

Mauger, D.M., Cabral, B.J., Presnyak, V., Su, S.V., Reid, D.W., Goodman, B., Link, K., Khatwani, N., Reynders, J., Moore, M.J., McFadyen, I.J., 2019. mRNA structure regulates protein expression through changes in functional half-life. Proc. Natl. Acad. Sci. 116, 24075– 24083. 10.1073/pnas.1908052116

McConnell, B.B., Yang, V.W., 2010. Mammalian Krüppel-Like Factors in Health and Diseases. Physiol. Rev. 90, 1337–1381. 10.1152/physrev.00058.2009

Miranda, K.C., Huynh, T., Tay, Y., Ang, Y.-S., Tam, W.-L., Thomson, A.M., Lim, B., Rigoutsos, I., 2006. A Pattern-Based Method for the Identification of MicroRNA Binding Sites and Their Corresponding Heteroduplexes. Cell 126, 1203–1217. 10.1016/j.cell.2006.07.031

Nishikura, K., 2016. A-to-I editing of coding and non-coding RNAs by ADARs. Nat. Rev. Mol. Cell Biol. 17, 83–96. 10.1038/nrm.2015.4

Nudel, R., Thompson, W.K., Børglum, A.D., Hougaard, D.M., Mortensen, P.B., Werge, T., Nordentoft, M., Benros, M.E., 2022. Maternal pregnancy-related infections and autism spectrum disorder—the genetic perspective. Transl. Psychiatry 12, 334. 10.1038/s41398-022-02068-9

Park, E., Jiang, Y., Hao, L., Hui, J., Xing, Y., 2021. Genetic variation and microRNA targeting of A- to-I RNA editing fine tune human tissue transcriptomes. Genome Biol. 22, 77. 10.1186/s13059-021-02287-1

Patil, I., 2021. Visualizations with statistical details: The “ggstatsplot” approach. J. Open Source Softw. 6, 3167. 10.21105/joss.03167

Perez, G., Barber, G.P., Benet-Pages, A., Casper, J., Clawson, H., Diekhans, M., Fischer, C., Gonzalez, J.N., Hinrichs, A.S., Lee, C.M., Nassar, L.R., Raney, B.J., Speir, M.L., van Baren, M.J., Vaske, C.J., Haussler, D., Kent, W.J., Haeussler, M., 2024. The UCSC Genome Browser database: 2025 update. Nucleic Acids Res. 53, D1243–D1249. 10.1093/nar/gkae974

Picardi, E., D’Erchia, A.M., Lo Giudice, C., Pesole, G., 2017. REDIportal: a comprehensive database of A-to-I RNA editing events in humans. Nucleic Acids Res. 45, D750–D757. 10.1093/nar/gkw767

Pinto, Y., Cohen, H.Y., Levanon, E.Y., 2014. Mammalian conserved ADAR targets comprise only a small fragment of the human editosome. Genome Biol. 15, R5. 10.1186/gb-2014-15-1-r5

Piontkivska, H., Plonski, N., Miyamoto, M.M., Wayne, M.L., 2019. Explaining Pathogenicity of Congenital Zika and Guillain–Barré Syndromes: Does Dysregulation of RNA Editing Play a Role? BioEssays 41, 1800239. 10.1002/bies.201800239

Plonski, N.-M., Johnson, E., Frederick, M., Mercer, H., Fraizer, G., Meindl, R., Casadesus, G., Piontkivska, H., 2020. Automated Isoform Diversity Detector (AIDD): a pipeline for investigating transcriptome diversity of RNA-seq data. BMC Bioinformatics 21, 578. 10.1186/s12859-020-03888-6

Provenzano, G., Corradi, Z., Monsorno, K., Fedrizzi, T., Ricceri, L., Scattoni, M.L., Bozzi, Y., 2016. Comparative Gene Expression Analysis of Two Mouse Models of Autism: Transcriptome Profiling of the BTBR and En2−/− Hippocampus. Front. Neurosci. 10. 10.3389/fnins.2016.00396

R Core Team, 2023. R: A Language and Environment for Statistical Computing. R Foundation for Statistical Computing, Vienna, Austria.

Richards, S., Aziz, N., Bale, S., Bick, D., Das, S., Gastier-Foster, J., Grody, W.W., Hegde, M., Lyon, E., Spector, E., Voelkerding, K., Rehm, H.L., 2015. Standards and guidelines for the interpretation of sequence variants: a joint consensus recommendation of the American College of Medical Genetics and Genomics and the Association for Molecular Pathology. Genet. Med. 17, 405–424. 10.1038/gim.2015.30

Roberts, J.T., Patterson, D.G., King, V.M., Amin, S.V., Polska, C.J., Houserova, D., Crucello, A., Barnhill, E.C., Miller, M.M., Sherman, T.D., Borchert, G.M., 2018. ADAR Mediated RNA Editing Modulates MicroRNA Targeting in Human Breast Cancer. Process. Basel Switz. 6, 42. 10.3390/pr6050042

Rubenstein, J.L.R., Merzenich, M.M., 2003. Model of autism: increased ratio of excitation/inhibition in key neural systems. Genes Brain Behav. 2, 255–267. 10.1034/j.1601-183X.2003.00037.x

Rylaarsdam, L., Guemez-Gamboa, A., 2019. Genetic Causes and Modifiers of Autism Spectrum Disorder. Front. Cell. Neurosci. 13, 385. 10.3389/fncel.2019.00385

Sabourin, K.R., Reynolds, A., Schendel, D., Rosenberg, S., Croen, L.A., Pinto-Martin, J.A., Schieve, L.A., Newschaffer, C., Lee, L.-C., DiGuiseppi, C., 2019. Infections in Children with Autism Spectrum Disorder: Study to Explore Early Development (SEED). Autism Res. Off. J. Int. Soc. Autism Res. 12, 136–146. 10.1002/aur.2012

Sclabassi, E., Peret, S., Qian, C., Gao, Y., 2025. Pharmacological Interventions in Autism Spectrum Disorder: A Comprehensive Review of Mechanisms and Efficacy. Biomedicines 13, 3025. 10.3390/biomedicines13123025

Sherry, S.T., Ward, M.-H., Kholodov, M., Baker, J., Phan, L., Smigielski, E.M., Sirotkin, K., 2001. dbSNP: the NCBI database of genetic variation. Nucleic Acids Res. 29, 308–311. 10.1093/nar/29.1.308

Slotkin, W., Nishikura, K., 2013. Adenosine-to-inosine RNA editing and human disease. Genome Med. 5, 105. 10.1186/gm508

Solomon, O., Di Segni, A., Cesarkas, K., Porath, H.T., Marcu-Malina, V., Mizrahi, O., Stern-Ginossar, N., Kol, N., Farage-Barhom, S., Glick-Saar, E., Lerenthal, Y., Levanon, E.Y., Amariglio, N., Unger, R., Goldstein, I., Eyal, E., Rechavi, G., 2017. RNA editing by ADAR1 leads to context-dependent transcriptome-wide changes in RNA secondary structure. Nat. Commun. 8, 1440. 10.1038/s41467-017-01458-8

Solomon, O., Oren, S., Safran, M., Deshet-Unger, N., Akiva, P., Jacob-Hirsch, J., Cesarkas, K., Kabesa, R., Amariglio, N., Unger, R., Rechavi, G., Eyal, E., 2013. Global regulation of alternative splicing by adenosine deaminase acting on RNA (ADAR). RNA 19, 591–604. 10.1261/rna.038042.112

Song, N., Nakagawa, S., Izumi, T., Toda, H., Kato, A., Boku, S., Inoue, T., Sakagami, H., Li, X., Koyama, T., 2013. Involvement of CaMKIV in neurogenic effect with chronic fluoxetine treatment. Int. J. Neuropsychopharmacol. 16, 803–812. 10.1017/S1461145712000570

Stahl, S.M., 1998. Mechanism of action of serotonin selective reuptake inhibitors: Serotonin receptors and pathways mediate therapeutic effects and side effects. J. Affect. Disord. 51, 215–235. 10.1016/S0165-0327(98)00221-3

Syafruddin, S.E., Mohtar, M.A., Wan Mohamad Nazarie, W.F., Low, T.Y., 2020. Two Sides of the Same Coin: The Roles of KLF6 in Physiology and Pathophysiology. Biomolecules 10, 1378. 10.3390/biom10101378

Sztainberg, Y., Zoghbi, H.Y., 2016. Lessons learned from studying syndromic autism spectrum disorders. Nat. Neurosci. 19, 1408–1417. 10.1038/nn.4420

Tan, M.H., Li, Q., Shanmugam, R., Piskol, R., Kohler, J., Young, A.N., Liu, K.I., Zhang, R., Ramaswami, G., Ariyoshi, K., Gupte, A., Keegan, L.P., George, C.X., Ramu, A., Huang, N., Pollina, E.A., Leeman, D.S., Rustighi, A., Goh, Y.P.S., Chawla, A., Del Sal, G., Peltz, G., Brunet, A., Conrad, D.F., Samuel, C.E., O’Connell, M.A., Walkley, C.R., Nishikura, K., Li, J.B., 2017. Dynamic landscape and regulation of RNA editing in mammals. Nature 550, 249–254. 10.1038/nature24041

Tanaka, H., Grooms, S.Y., Bennett, M.V.L., Zukin, R.S., 2000. The AMPAR subunit GluR2: still front and center-stage1. Brain Res., Towards 2010, A brain Odyssey, The 3rd Brain Research Interactive 886, 190–207. 10.1016/S0006-8993(00)02951-6

Tariq, A., Piontkivska, H., 2024. Reovirus infection induces transcriptome-wide unique A-to-I editing changes in the murine fibroblasts. Virus Res. 346, 199413. 10.1016/j.virusres.2024.199413

Tartaglione, A.M., Villani, A., Ajmone-Cat, M.A., Minghetti, L., Ricceri, L., Pazienza, V., De Simone, R., Calamandrei, G., 2022. Maternal immune activation induces autism-like changes in behavior, neuroinflammatory profile and gut microbiota in mouse offspring of both sexes. Transl. Psychiatry 12, 384. 10.1038/s41398-022-02149-9

Tate, K., Kirk, B., Tseng, A., Ulffers, A., Litwa, K., 2021. Effects of the Selective Serotonin Reuptake Inhibitor Fluoxetine on Developing Neural Circuits in a Model of the Human Fetal Cortex. Int. J. Mol. Sci. 22, 10457. 10.3390/ijms221910457

Than, U.T.T., Nguyen, L.T., Nguyen, P.H., Nguyen, X.-H., Trinh, D.P., Hoang, D.H., Nguyen, P.A.T., Dang, V.D., 2023. Inflammatory mediators drive neuroinflammation in autism spectrum disorder and cerebral palsy. Sci. Rep. 13, 22587. 10.1038/s41598-023-49902-8

Tiraboschi, E., Tardito, D., Kasahara, J., Moraschi, S., Pruneri, P., Gennarelli, M., Racagni, G., Popoli, M., 2004. Selective Phosphorylation of Nuclear CREB by Fluoxetine is Linked to Activation of CaM Kinase IV and MAP Kinase Cascades. Neuropsychopharmacology 29, 1831–1840. 10.1038/sj.npp.1300488

Tomaselli, S., Bonamassa, B., Alisi, A., Nobili, V., Locatelli, F., Gallo, A., 2013. ADAR Enzyme and miRNA Story: A Nucleotide that Can Make the Difference. Int. J. Mol. Sci. 14, 22796–22816. 10.3390/ijms141122796

Tran, S.S., Jun, H.-I., Bahn, J.H., Azghadi, A., Ramaswami, G., Van Nostrand, E.L., Nguyen, T.B., Hsiao, Y.-H.E., Lee, C., Pratt, G.A., Martínez-Cerdeño, V., Hagerman, R.J., Yeo, G.W., Geschwind, D.H., Xiao, X., 2019. Widespread RNA editing dysregulation in brains from autistic individuals. Nat. Neurosci. 22, 25–36. 10.1038/s41593-018-0287-x

Tsivion-Visbord, H., Kopel, E., Feiglin, A., Sofer, T., Barzilay, R., Ben-Zur, T., Yaron, O., Offen, D., Levanon, E.Y., 2020. Increased RNA editing in maternal immune activation model of neurodevelopmental disease. Nat. Commun. 11, 5236. 10.1038/s41467-020-19048-6

Valinezhad Orang, A., Safaralizadeh, R., Kazemzadeh-Bavili, M., 2014. Mechanisms of miRNA-Mediated Gene Regulation from Common Downregulation to mRNA-Specific Upregulation. Int. J. Genomics 2014, 970607. 10.1155/2014/970607

Wales-McGrath, B., Mercer, H., Piontkivska, H., 2023. Changes in ADAR RNA editing patterns in CMV and ZIKV congenital infections. BMC Genomics 24, 685. 10.1186/s12864-023-09778-4

Walkley, C.R., Li, J.B., 2017. Rewriting the transcriptome: adenosine-to-inosine RNA editing by ADARs. Genome Biol. 18, 205. 10.1186/s13059-017-1347-3

Wang, I.X., So, E., Devlin, J.L., Zhao, Y., Wu, M., Cheung, V.G., 2013. ADAR Regulates RNA Editing, Transcript Stability, and Gene Expression. Cell Rep. 5, 849–860. 10.1016/j.celrep.2013.10.002

Wang, L., Wang, B., Wu, C., Wang, J., Sun, M., 2023. Autism Spectrum Disorder: Neurodevelopmental Risk Factors, Biological Mechanism, and Precision Therapy. Int. J. Mol. Sci. 24, 1819. 10.3390/ijms24031819

Wang, Y., Li, J.-T., Zhu, L.-L., Wu, Y.-K., Su, Y.-A., Si, T.-M., 2026. Mitochondrial-inflammation crosstalk in major depressive disorder: molecular mechanisms and therapeutic implications. Mol. Psychiatry 1–12. 10.1038/s41380-026-03732-y

Wang, Y., Neumann, M., Hansen, K., Hong, S.M., Kim, S., Noble-Haeusslein, L.J., Liu, J., 2011. Fluoxetine Increases Hippocampal Neurogenesis and Induces Epigenetic Factors But Does Not Improve Functional Recovery after Traumatic Brain Injury. J. Neurotrauma 28, 259–268. 10.1089/neu.2010.1648

Weissmann, D., Van Der Laan, S., Underwood, M.D., Salvetat, N., Cavarec, L., Vincent, L., Molina, F., Mann, J.J., Arango, V., Pujol, J.F., 2016. Region-specific alterations of A-to-I RNA editing of serotonin 2c receptor in the cortex of suicides with major depression. Transl. Psychiatry 6, e878–e878. 10.1038/tp.2016.121

Wong, A., Zhou, A., Cao, X., Mahaganapathy, V., Azaro, M., Gwin, C., Wilson, S., Buyske, S., Bartlett, C.W., Flax, J.F., Brzustowicz, L.M., Xing, J., 2022. MicroRNA and MicroRNA-Target Variants Associated with Autism Spectrum Disorder and Related Disorders. Genes 13, 1329. 10.3390/genes13081329

Wong, D.T., Horng, J.S., Bymaster, F.P., Hauser, K.L., Molloy, B.B., 1974. A selective inhibitor of serotonin uptake: Lilly 110140, 3-(p-Trifluoromethylphenoxy)-n-methyl-3-phenylpropylamine. Life Sci. 15, 471–479. 10.1016/0024-3205(74)90345-2

Wong, D.T., Perry, K.W., Bymaster, F.P., 2005. The Discovery of Fluoxetine Hydrochloride (Prozac). Nat. Rev. Drug Discov. 4, 764–774. 10.1038/nrd1821

Wright, A.L., Konen, L.M., Mockett, B.G., Morris, G.P., Singh, A., Burbano, L.E., Milham, L., Hoang, M., Zinn, R., Chesworth, R., Tan, R.P., Royle, G.A., Clark, I., Petrou, S., Abraham, W.C., Vissel, B., 2023. The Q/R editing site of AMPA receptor GluA2 subunit acts as an epigenetic switch regulating dendritic spines, neurodegeneration and cognitive deficits in Alzheimer’s disease. Mol. Neurodegener. 18, 65. 10.1186/s13024-023-00632-5

Wu, T., Hu, E., Xu, S., Chen, M., Guo, P., Dai, Z., Feng, T., Zhou, L., Tang, W., Zhan, L., Fu, X., Liu, S., Bo, X., Yu, G., 2021. clusterProfiler 4.0: A universal enrichment tool for interpreting omics data. The Innovation 2, 100141. 10.1016/j.xinn.2021.100141

Yang, Y., Okada, S., Sakurai, M., 2021. Adenosine-to-inosine RNA editing in neurological development and disease. RNA Biol. 18, 999–1013. 10.1080/15476286.2020.1867797

Yang, Y., Zhou, X., Jin, Y., 2013. ADAR-mediated RNA editing in non-coding RNA sequences. Sci. China Life Sci. 56, 944–952. 10.1007/s11427-013-4546-5

Yuan, J., Xu, L., Bao, H.-J., Wang, J., Zhao, Y., Chen, S., 2023. Biological roles of A-to-I editing: implications in innate immunity, cell death, and cancer immunotherapy. J. Exp. Clin. Cancer Res. 42, 149. 10.1186/s13046-023-02727-9

Yue, F., Cheng, Y., Breschi, A., Vierstra, J., Wu, W., Ryba, T., Sandstrom, R., Ma, Z., Davis, C., Pope, B.D., Shen, Y., Pervouchine, D.D., Djebali, S., Thurman, R.E., Kaul, R., Rynes, E., Kirilusha, A., Marinov, G.K., Williams, B.A., Trout, D., Amrhein, H., Fisher-Aylor, K., Antoshechkin, I., DeSalvo, G., See, L.-H., Fastuca, M., Drenkow, J., Zaleski, C., Dobin, A., Prieto, P., Lagarde, J., Bussotti, G., Tanzer, A., Denas, O., Li, K., Bender, M.A., Zhang, M., Byron, R., Groudine, M.T., McCleary, D., Pham, L., Ye, Z., Kuan, S., Edsall, L., Wu, Y.-C., Rasmussen, M.D., Bansal, M.S., Kellis, M., Keller, C.A., Morrissey, C.S., Mishra, T., Jain, D., Dogan, N., Harris, R.S., Cayting, P., Kawli, T., Boyle, A.P., Euskirchen, G., Kundaje, A., Lin, S., Lin, Y., Jansen, C., Malladi, V.S., Cline, M.S., Erickson, D.T., Kirkup, V.M., Learned, K., Sloan, C.A., Rosenbloom, K.R., Lacerda de Sousa, B., Beal, K., Pignatelli, M., Flicek, P., Lian, J., Kahveci, T., Lee, D., James Kent, W., Ramalho Santos, M., Herrero, J., Notredame, C., Johnson, A., Vong, S., Lee, K., Bates, D., Neri, F., Diegel, M., Canfield, T., Sabo, P.J., Wilken, M.S., Reh, T.A., Giste, E., Shafer, A., Kutyavin, T., Haugen, E., Dunn, D., Reynolds, A.P., Neph, S., Humbert, R., Scott Hansen, R., De Bruijn, M., Selleri, L., Rudensky, A., Josefowicz, S., Samstein, R., Eichler, E.E., Orkin, S.H., Levasseur, D., Papayannopoulou, T., Chang, K.-H., Skoultchi, A., Gosh, S., Disteche, C., Treuting, P., Wang, Y., Weiss, M.J., Blobel, G.A., Cao, X., Zhong, S., Wang, T., Good, P.J., Lowdon, R.F., Adams, L.B., Zhou, X.-Q., Pazin, M.J., Feingold, E.A., Wold, B., Taylor, J., Mortazavi, A., Weissman, S.M., Stamatoyannopoulos, J.A., Snyder, M.P., Guigo, R., Gingeras, T.R., Gilbert, D.M., Hardison, R.C., Beer, M.A., Ren, B., 2014. A comparative encyclopedia of DNA elements in the mouse genome. Nature 515, 355–364. 10.1038/nature13992

Zaidan, H., Ramaswami, G., Barak, M., Li, J.B., Gaisler-Salomon, I., 2018. Pre-reproductive stress and fluoxetine treatment in rats affect offspring A-to-I RNA editing, gene expression and social behavior. Environ. Epigenetics 4, dvy021. 10.1093/eep/dvy021

Zhang, Y., You, X., Li, S., Long, Q., Zhu, Y., Teng, Z., Zeng, Y., 2020. Peripheral Blood Leukocyte RNA-Seq Identifies a Set of Genes Related to Abnormal Psychomotor Behavior Characteristics in Patients with Schizophrenia. Med. Sci. Monit. Int. Med. J. Exp. Clin. Res. 26, e922426-1–e922426-31. 10.12659/MSM.922426

Zhu, T., Su, Q., Wang, C., Shen, L., Chen, H., Feng, S., Peng, X., Chen, S., Wang, Y., Jiang, H., Chen, J., 2021. SDF4 Is a Prognostic Factor for 28-Days Mortality in Patients With Sepsis via Negatively Regulating ER Stress. Front. Immunol. 12. 10.3389/fimmu.2021.659193

